# Fragmented social networks promote complex behavioural contagions over infectious disease spread

**DOI:** 10.1101/2025.08.28.672836

**Authors:** Nitara Wijayatilake, Shweta Bansal, Amy B Pederson, Matthew Silk

## Abstract

Group living and social interactions among animals provide key benefits, such as the exchange of beneficial social information and novel behaviours, but also pose the risk of spreading costly infectious diseases, presenting a social trade-off. While both information and infections spread across social networks, they typically have distinct mechanisms of transmission. Here, we model social information and behaviour spread as a complex contagion governed by a conformist learning rule, while pathogen transmission follows a simple contagion mechanism. Building on theoretical foundations, our study applies computational models to examine how subgroup structure (modularity) influences the spread of these contagions across diverse animal social structures sourced from the Animal Social Network Repository (ASNR). Our findings reveal that high modularity and subgroup structure slow simple contagion spread, whereas complex contagions are less impeded by this fragmentation. Consequently, our results suggest that social networks divided into small groups or subgroups can help balance the competing pressures of acquiring social information and avoiding infectious disease in real-world networks.

## Introduction

Animal social systems exhibit remarkable diversity, from largely solitary species to those forming stable groups and differentiated social bonds, with many intermediate forms of social system and intraspecific variation in interaction patterns [1, 2, 3]. Social interactions with conspecifics can offer significant advantages, including social transmission, defined here as the spread of information and behaviours [4, 5]. Through social learning, individuals can acquire novel behaviours from their peers related to habitat selection, resource acquisition, and anti-predator strategies, which can enhance their fitness [6, 7, 8]. However, sociality comes with costs, such as increasing the risk of pathogen and parasite transmission [9, 10].

Individuals that live in groups face both the advantages and risks of social interactions. Dolphins, for example, exhibit extensive social learning behaviours related to foraging, hunting, and communication. Young dolphins learn complex hunting techniques, such as ‘sponging’ —a foraging behaviour involving sponges as tools—by observing their mothers [11]. However, their close social interactions also facilitate the spread of pathogens such as morbillivirus [12]. The simultaneous yet opposing forces of pathogen transmission and social transmission represent a social trade-off [13, 14]. Effectively balancing access and use of beneficial social information with the avoidance of harmful infections is therefore likely to promote individual fitness [15, 16, 17] and potentially group persistence. Despite this, we still understand little about how different animal social structures jointly influence social contagions and pathogen spread, making it difficult to establish the role of this trade-off in the evolutionary ecology of animal social systems.

The concept of a ‘contagion’ is commonly employed to describe the spread of diseases or ideas from one individual to another. While both pathogens and behaviour are frequently modelled as contagions, the mechanisms governing their spread can differ substantially. The spread of a pathogen to a susceptible individual for a directly-transmitted infection typically depends on increased exposure to infected individuals, with each interaction contributing independently to the likelihood of transmission [18]. In contrast, the acquisition and use of social information often involves a more complex process, where the effect of each interaction is not independent but instead shaped by cumulative or reinforcing dynamics. Complex behaviours like tool-use or mate-choice may require multiple exposures or social reinforcement to spread effectively to new individuals [8, 19, 20]. For this reason, social contagions are often modelled as complex contagions, where multiple exposures are required before transmission occurs [21, 22]. In contrast, directly transmitted infectious diseases are typically treated as simple contagions, where each contact with an infected individual carries an independent probability of spread [23, 24]. This distinction aids in effectively modelling the differing dynamics of social contagions versus pathogen spread as it accounts for nuances inherent to each transmission process [25, 26]. With these differences in transmission formalised, it becomes important to examine how key factors, namely social structure, can influence and differentiate the spread of disease and social contagions [22].

Animals may employ various learning strategies, such as copying the behaviour of familiar individuals or conforming to majority behaviours, to uptake social contagions within their groups [8]. One proposed rule is conformist learning, where the prevalence of the behaviour is what guides the adoption process, with an individual more likely to imitate a behaviour when the majority of their neighbours have already adopted it [27, 28]. In animal behaviour and sociology, this is colloquially termed ‘copy the majority’, and it is widely seen in the human and animal world [29, 30]. Research in great tits, *Parus major*, found that social learning is preferred over asocial learning, with most individuals choosing to adopt the behaviour demonstrated by the majority of their social contacts [31]. In a different study, nine-spined sticklebacks exhibited a disproportionately strong preference for majority behaviour when choosing feeding sites, indicating a high level of conformist learning [32]. When social contagions spread through conformist learning, differences in social network structure could illuminate how groups balance the competing pressures of social information and infectious disease [16].

Many animal social networks are subdivided into discrete social communities (e.g. based on groups or subgroups)[33, 34]. Here, we refer to communities as densely connected subgroups with more connections within than between groups [35], and modularity as a measure of the strength of such subdivision in a network [36]. Within communities, the high density of connections can intensify the spread of contagions [37]. Community structure can influence both disease [38, 39, 40] and social [41] contagions. Sah et al. [40] found that infection burden was lowest in social networks with high modularity and fragmentation, where subgroups acted as structural barriers. However, independently of community size, these effects only became pronounced at very high modularity levels. Interestingly, due to social reinforcement, complex contagions governed by threshold dynamics have been shown to spread most effectively at intermediate levels of modularity, which balance dense intra-community ties with sufficient inter-community connectivity [41]. Understanding how social structures mitigate selection pressures—minimising disease transmission while enhancing the spread of beneficial information—could offer valuable insights into how these ecological dynamics contribute to the evolution of group and sub-group structure in animal social systems.

Evans et al. [42] revealed that highly modular networks with small groups favoured the spread of a conformist over a simple contagion, suggesting that networks with numerous, small subgroups might regulate the trade-offs between social and infection transmission within the same set of social interactions. However, the idea that modular network structures may help balance the trade-off between infection and information has not been tested in real-world systems. This gap restricts our understanding of how natural social systems balance competing selection pressures on sociality. To address this, we draw on the unique breadth of data offered by the Animal Social Network Repository (ASNR) to compare the speed of spread for both simple and complex contagions within diverse real-world networks [43]. We expect that community-structured networks with high modularity will slow the spread of both contagions, with this effect being most pronounced when the number of communities is highest (i.e. sub-groups are smaller). Additionally, we expect (complex) social contagions to navigate modular structures more effectively than (simple) infectious contagions, resulting in their speed of spread being less impeded by community structure. Simulating contagion spread on real-world networks offers critical insight into how realistic social structures influence the interplay between pathogen and information transmission, capturing dynamics that are difficult to observe in purely theoretical models. Therefore, by modelling contagions through empirical animal social networks, our study bridges the gap between theoretical and real-world structures, helping to quantify the potential for animal social group structure to influence the ecological consequences of social interactions.

## Methods

All analyses were performed using R version 4.4.2 [44]. All code is available at our Zenodo repository.

### Animal Social Network Repository (ASNR)

Empirical networks were accessed from the ASNR version 3.0 [43, 45], an open-source mixed species data repository of animal social behaviour networks spanning several taxonomic groups including insects, fish, birds, reptiles, and mammals. We selected networks from the top three quartiles of the node count distribution in the ASNR to ensure the networks were large enough to support contagion dynamics and subgroup formation, reducing our dataset from 1,030 to 712. Next, we restricted our analysis to weighted networks, yielding a final dataset of 545 networks spanning 52 species and exhibiting a wide range of fragmentation (number of sub-groups) and modularity values (Figure 1). Supplementary Figure 1 shows that the networks span a representative range of characteristics across ASNR. Simulations were performed on the largest connected component of each network, ensuring that contagions could feasibly reach all individuals.

**Figure 1:**
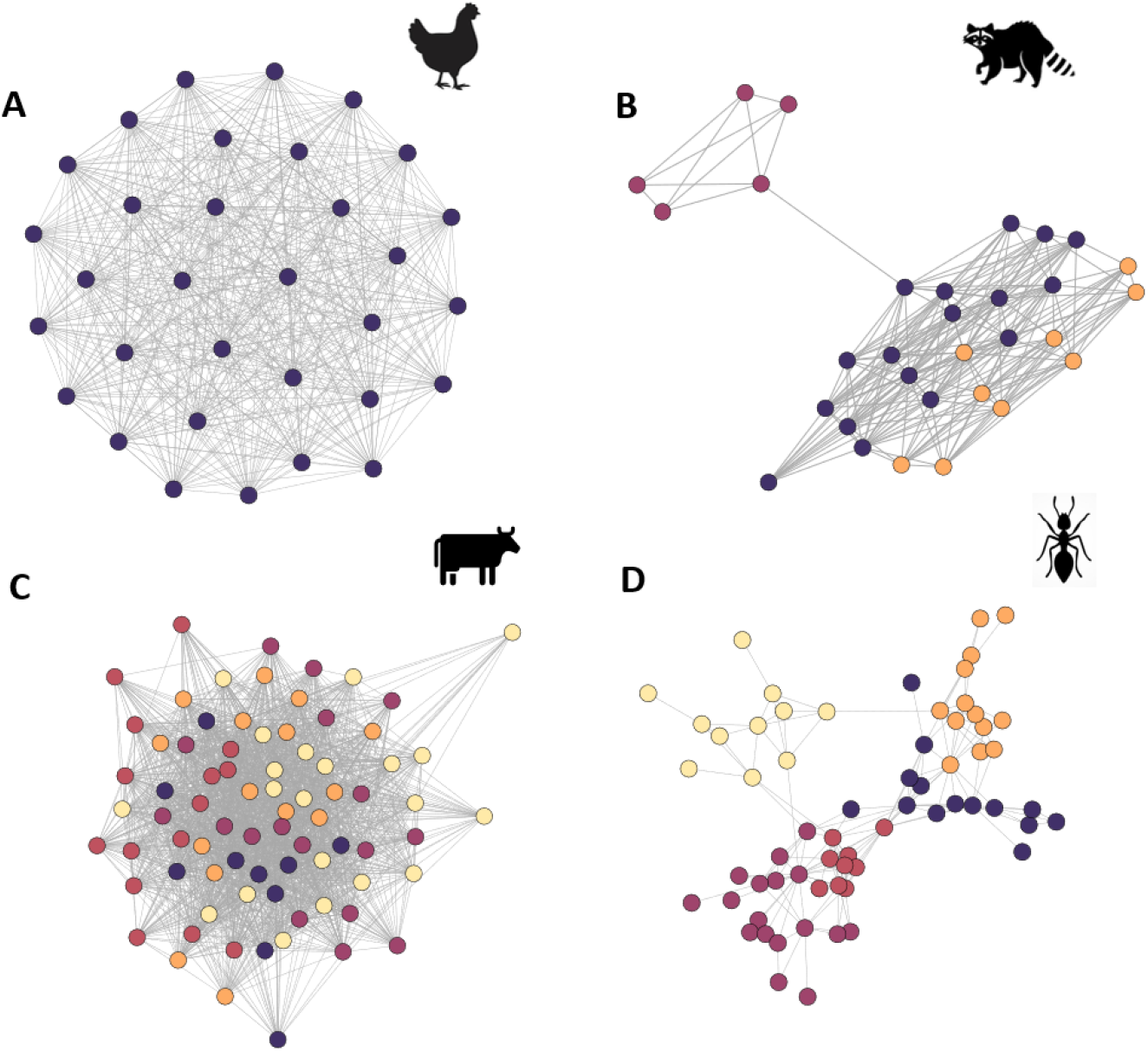
Example networks from the Animal Social Network Repository (ASNR), demonstrating variations in modularity and community structure. (A) *Gallus gallus domesticus* (chicken): modularity = 0, single community. (B) *Procyon lotor* (raccoon): modularity = 0.4, three communities. (C) *Bos taurus* (cow): modularity = 0.1, five communities. (D) *Camponotus pennsylvanicus* (ant): modularity = 0.5, five communities.

### Contagion spread

A susceptible-infected (SI) model was used to simulate the spread of both simple and complex contagions. The model focused solely on the transmission phase, to capture contagion spread dynamics and directly compare the initial uptake of simple and complex contagions. Baseline networks were generated using Erdős–Rényi random graphs, designed to match the size (number of nodes) and edge density of each empirical network. These baseline networks served as controls to ensure comparable conditions for the transmissibility of both contagions. The simulation was seeded at a random node in the network and allowed to spread through the network with a given transmission probability of a node changing from susceptible (uninfected) to infected status. Since simulations used weighted networks, stronger associations increased the likelihood of contagion spread.

For the simple contagion (pathogen) transmission, for each time step, the probability of infection for a node *v* is:

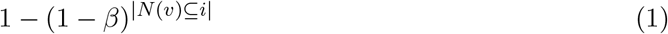

where *β* is the transmission rate and |*N* (*v*) ⊆ *i*| is the weighted sum of infected neighbours. In our SI model, R_0_ is defined as the per-time period basic reproduction number—that is, the average number of secondary infections produced by an infected individual in a fully susceptible network over a fixed time interval. We calculate the per-time-step transmission probability *β* from *R*_0_ and the network’s degree distribution as:

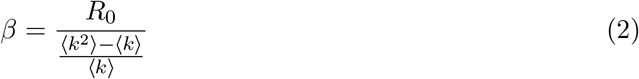

where ⟨*k*⟩ is the average degree and ⟨*k*^2^⟩ is the average squared degree. This adjusts for how the network’s connectivity influences transmission. An *R*_0_ of 1.5 was chosen based on findings from Evans et al. [25], which demonstrated that intermediate transmissibility values resulted in the greatest differences in transmission rates between social and disease transmission.

We model social transmission as a complex contagion using a conformist learning rule, where the likelihood an individual adopts information is determined by a sigmoidal function of the proportion of its neighbours that are informed, meaning that an individual only becomes more likely to adopt the social contagion when a high proportion of its contacts are informed (and not just because it has more contacts overall).

In each time-step, the probability of node *v* becoming informed is:

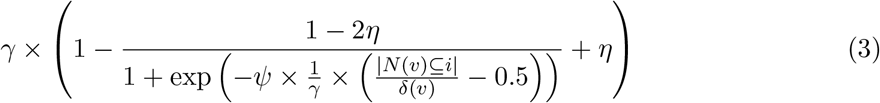

where *γ* is the scaling parameter to modify the rate of social contagion spread, |*N* (*v*) ⊆ *i*| is the weighted sum of informed neighbours, and *δ*(*v*) is the node’s degree, *η* is the base learning rate (to allow for a small amount of independent learning) and *ψ* is the steepness of the sigmoidal function of social contagion uptake [42]. After exploring the parameter space, we set *η* at 0.001 to allow minimal independent learning, while a *ψ* of 10 effectively captured the steep dynamics of conformist learning.

To simulate transmission through social networks, we adapted the method from Evans et al. [42]. We measure the time for 75% of the network to become infected to capture the speed of contagion spread through the majority of the network. To determine an appropriate value for *γ*, the spreading parameter governing the rate of social contagion, we first calibrated contagion dynamics within the baseline (random) networks. This calibration allowed us to select a *γ* value that ensured both pathogen and social contagions spread at equivalent rates under identical network conditions. Doing so was important because, when later moving simulations into the empirical structures, we could isolate and examine the effects of community structure on the differences in dynamics between the two contagion types, rather than attributing differences solely to transmissibility.

To achieve this, we began by running 50 simulations of pathogen spread on the baseline networks, recording the time required for 75% of individuals in the network to become infected. This established a benchmark for simple contagion dynamics. Next, social contagion spread was calibrated by testing a range of *γ* values, recording the time it took for 75% of individuals in the network to adopt the social contagion. We used the Bhattacharyya distance (a metric for quantifying the similarity between probability distributions) to evaluate the overlap between the spread time distributions of pathogen transmission and those of social contagion for each *γ* value [46]. This approach enabled us to identify the optimal *γ* value that most closely aligned the extent of social contagion spread with pathogen spread. Using the chosen optimal *γ* value, we then simulated the concurrent spread of infection and information across each empirical network. Each simulation was repeated 50 times per network, seeding in a randomly selected node each time, and we recorded the time required for 75% of the network nodes to become either infected with disease or informed with a social contagion. Figure 2 summarises the steps involved in our data collection and simulation processes.

**Figure 2:**
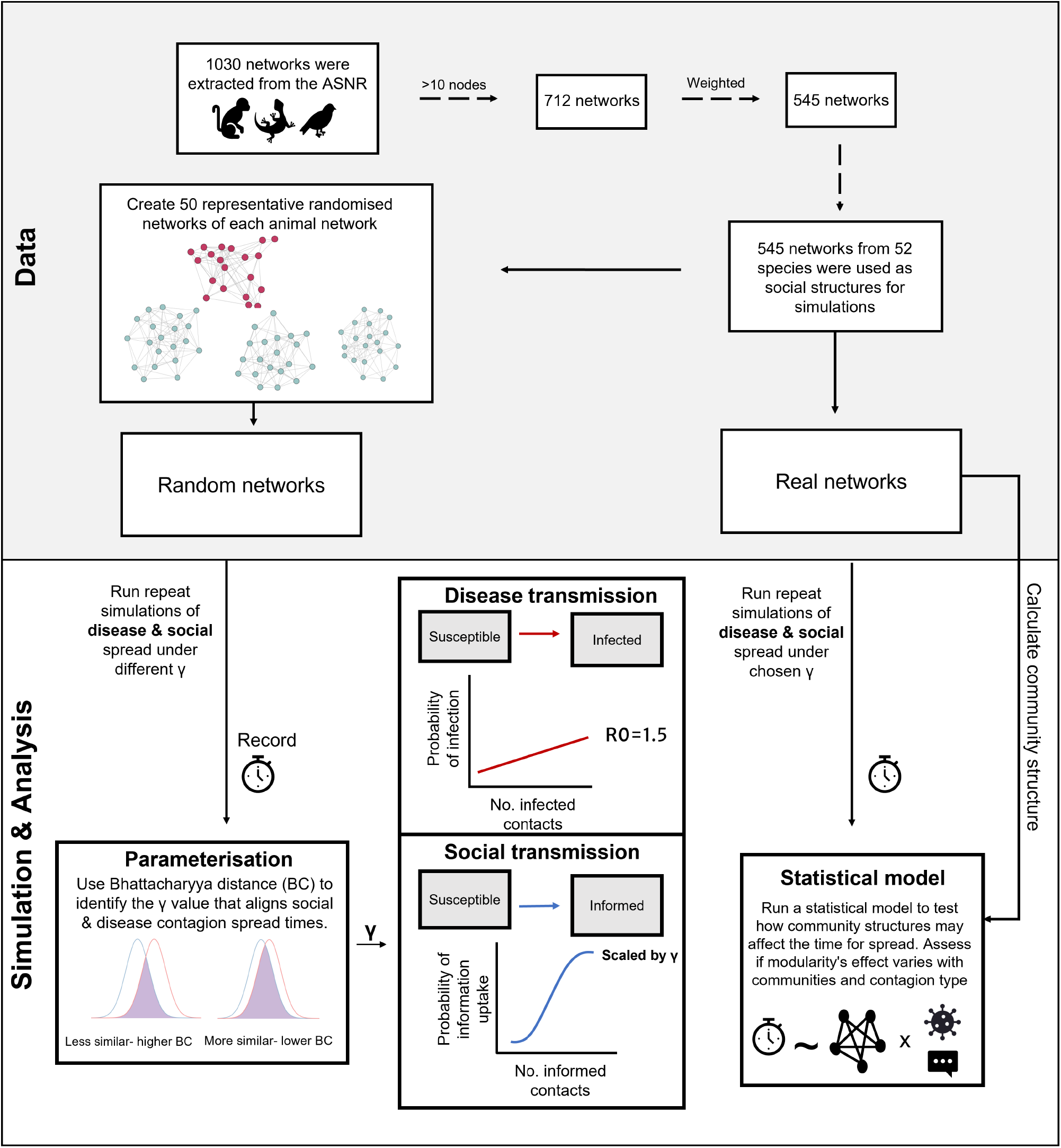
Overview of data extraction and simulation pipeline

### Network Metric Calculations

To investigate network structure as a predictor of relative contagion transmission differences, we calculated modularity, subgroup cohesion, network size (the number of nodes), edge density (proportion of possible edges present), and network fragmentation (the number of communities) of the empirical networks. Average module size and clustering coefficient were also calculated but excluded from analyses due to high collinearity (Pearson’s r *>* 0.7) with other variables (Supplementary Figure 2). These metrics were calculated using `igraph` [47]. Subgroup cohesion was measured as the proportion of total contacts that occur within subgroups [40]. For community detection, we employed a multi-level modularity optimisation algorithm using `cluster louvain` [35]. The number of communities served as an index of network fragmentation, reflecting both the number of subgroups and their relative sizes; as network size is controlled, a higher community count corresponds with smaller subgroup sizes. Modularity (*Q*) was calculated following Newman’s (2004) definition, which compares the density of edges within versus between communities [36], while *Q*_rel_ was computed using the formulation of Sah et al. [40] to account for variations in network size and density (Supplementary Information).

### Statistical model

We employed Bayesian regression models using the `brms` package in `R` to investigate how network structure influences the time taken for the contagion to reach 75% of the network for disease and social contagions [48]. The response variable was the time required for contagion spread to reach 75% of the population, with fixed effects for contagion type (pathogen vs. social), modularity, number of communities, network size, and edge density, as well as interactions among modularity, number of communities, and contagion type. All numeric predictors were centred to improve interpretability. To account for the hierarchical structure of the data, where multiple networks are nested within species, we included random intercepts for networks nested within species. This approach allows the model to capture variability in spread times both between species and among networks within species. We allowed the model’s residual standard deviation to vary across species and networks by modelling *σ* with random intercepts for networks nested within species; this accounts for potential heterogeneity in residual variance. We also ran a model with nested random effects for species within networks while assuming constant residual variance across groups. Model fit was assessed using Leave-One-Out Cross-Validation (LOO) [49] and the model that allowed residual variance to vary across species and networks was selected (Supplementary Information).

A skew-normal response distribution family was used to accommodate slight asymmetry in the tails of contagion spread times, providing a better fit than a symmetric Gaussian model. Posterior distributions were estimated using the default No-U-Turn Sampler (NUTS) algorithm, implemented across four Markov chains, each consisting of 2,000 iterations, with the initial 1,000 iterations designated for warmup. We conducted posterior predictive checks using density overlays to evaluate how well the model captured the observed data distribution (Supplementary Figure 4).

To assess robustness, we fit several supplementary models. One replaced Q_rel_ with subgroup cohesion to evaluate alternative measures of modular structure [40]. We also conducted two sensitivity analyses. Firstly, we excluded *Camponotus fellah* networks, which had a disproportionate number of observations (*n* = 221), to see if this species was driving the observed effect. Secondly, although we initially selected networks with at least 10 nodes for contagion simulations, visual inspection of the raw data revealed excessive variance in spread times for smaller and sparser networks. We therefore applied an additional quality filter, limited to networks with at least 20 nodes and density ≥ 0.15 (see Supplementary Information). In the main text, we present results from the full dataset (all 545 networks) to maximise sample size and ecological representation. Main findings remained consistent across all models (Supplementary Information).

## Results

The selected networks from the ASNR represent a wide diversity of network structures (Table 1). Approximately 45% of networks had Q_rel_ scores between 0.5 and 0.7 and between three and four communities. Figure 3 shows that our sample of real-world animal social networks span a broad range of variation in potential animal social structures. Most networks had either a low Q_rel_ and small number of communities or a high Q_rel_ and larger number of communities (higher fragmentation).

**Table 1:** Summary of ASNR Network Characteristics selected for our model

| Metric | Range | Central Tendency |
| --- | --- | --- |
| $Q_{rel}$ | 0.01 – 1.0 | Mean = 0.68 |
| Number of Communities | 1 – 12 | Median = 4 |
| Edge Density | 0.11 – 1.00 | Mean = 0.50 |
| Network Size | 10 – 252 | Median = 51 |

**Figure 3:**
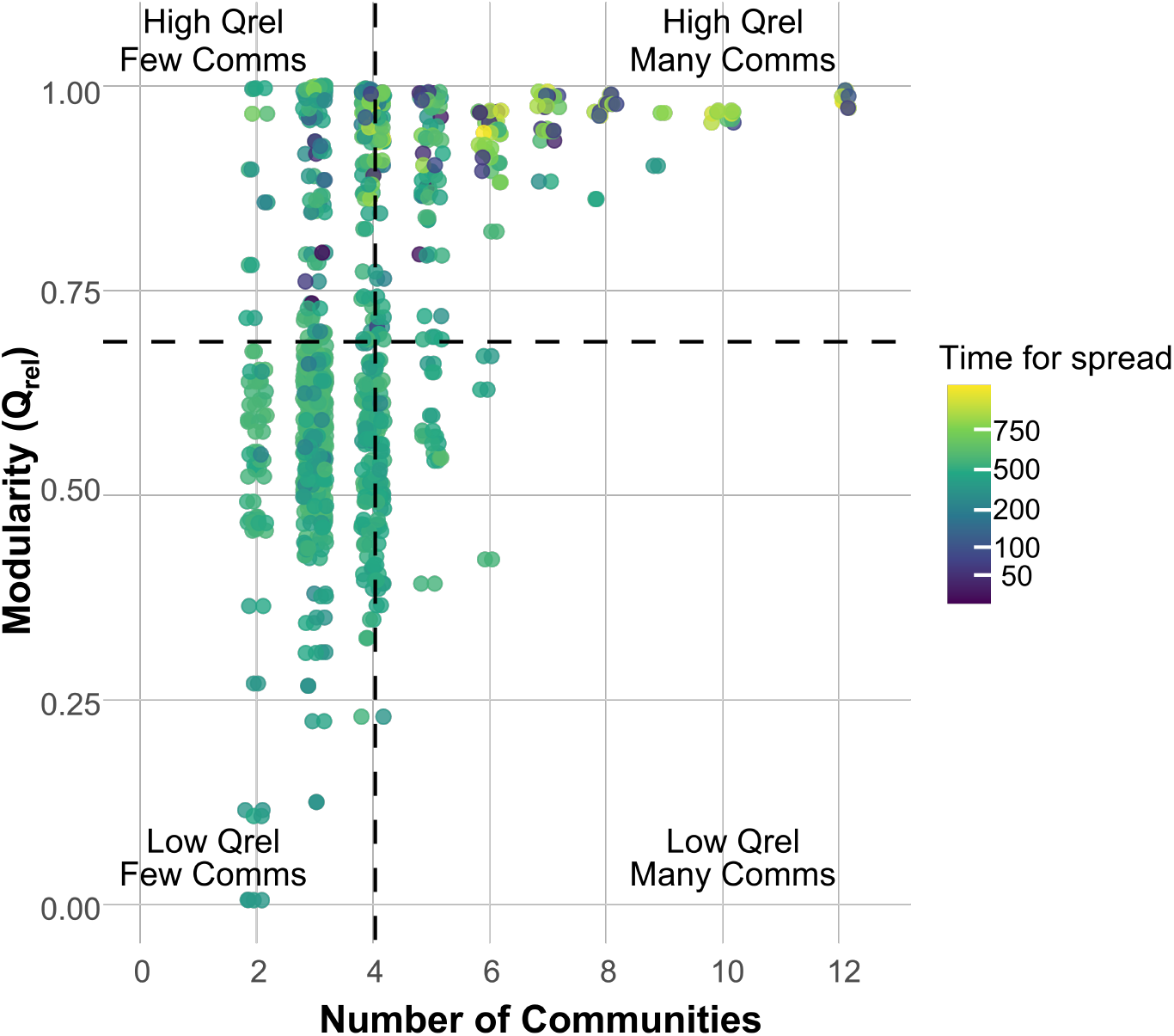
Relationship between modularity (Q_rel_) and number of communities across networks. Dashed lines indicate median values. Quadrant labels highlight combinations of (Q_rel_) and community number. Points are coloured by the time taken for contagion spread to reach 75% of the network. Networks with number of communities =1 are excluded, as Q_rel_ is not meaningful for a single community.

Our model results showed that networks with stronger community structures, indicated by higher modularity and more communities (higher fragmentation), took more time for 75% of the network to become infected by disease contagions. In contrast, social contagions reached 75% of the network in less time as network fragmentation increased, demonstrating that complex contagions more effectively traverse community boundaries than disease contagions.

The three-way interaction among contagion type, modularity, and number of communities (*β* = −98.31, 95% CI: −105.85, −90.70) indicated that it is the combination of modularity and network fragmentation (number of communities) that affects contagion dynamics in animal social networks. When networks had few communities, the spread of both disease and social contagions was unaffected by modularity. In contrast, in networks with many communities, social contagions were better able to overcome the combined effects of high modularity and fragmentation, while disease contagions were increasingly delayed. Social contagions even spread faster in more fragmented and modular networks (Figure 4). These findings indicate that modularity strongly influences contagion dynamics only when fragmentation is high.

**Figure 4:**
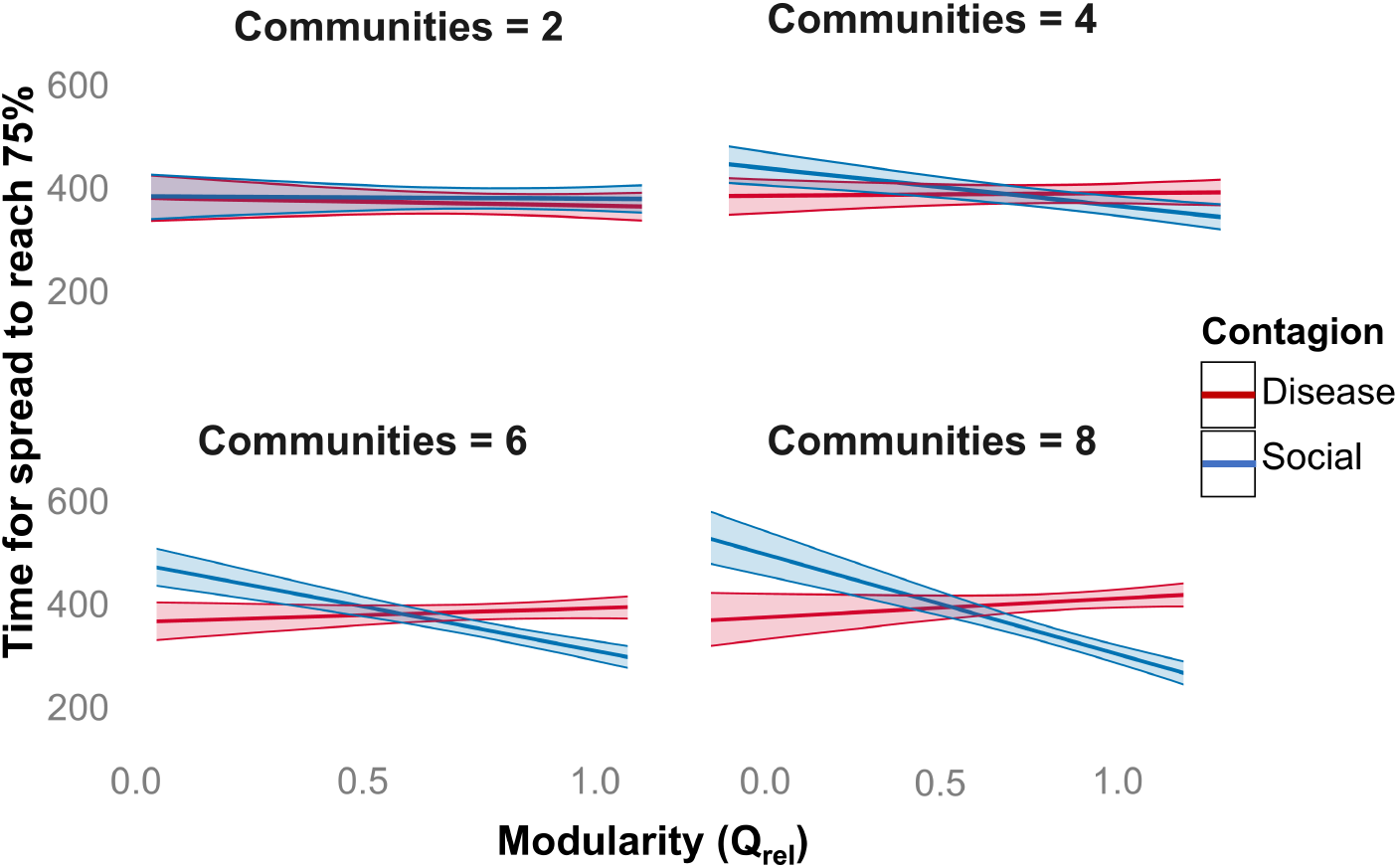
Influence of modularity (uncentred) on time to reach 75% spread across networks with varying numbers of communities (2, 4, 6, and 8) in disease contagions (red), and social contagions (blue)- the shaded areas depict 95% confidence intervals.

Examination of two-way interactions within the model revealed further differences between disease and social contagions. For disease contagions, for an average modularity, networks with more communities generally increased the time taken for the spread to reach 75% of the network (*β* = 16.73, 95% CI: 10.31, 23.47; Figure 5C), while modularity alone did not show a clear effect on spread time for the average level of network fragmentation (*β* = −52.45, 95% CI: −128.32, 23.19; Figure 5A). The two-way interaction between contagion type and number of communities (*β* = −17.06, 95% CI: −18.80, −15.33) indicates that, at the average modularity, increasing fragmentation slows the spread of disease contagions (reference category), while complex social contagions remain unaffected or may even spread slightly faster. In contrast, the interaction between modularity and contagion type (*β* = −12.44, 95% CI: −49.77, 24.29) has a wide credible interval overlapping zero, suggesting no clear difference between contagion types in their response to modularity at average levels of network fragmentation. As expected, larger network size was associated with increased spread time (*β* = 0.87, 95% CI: 0.71, 1.03; Figure 5D), whereas higher network density was associated with decreased spread time (*β* = −102.69, 95% CI: −141.80, −64.76; Figure 5B).

**Figure 5:**
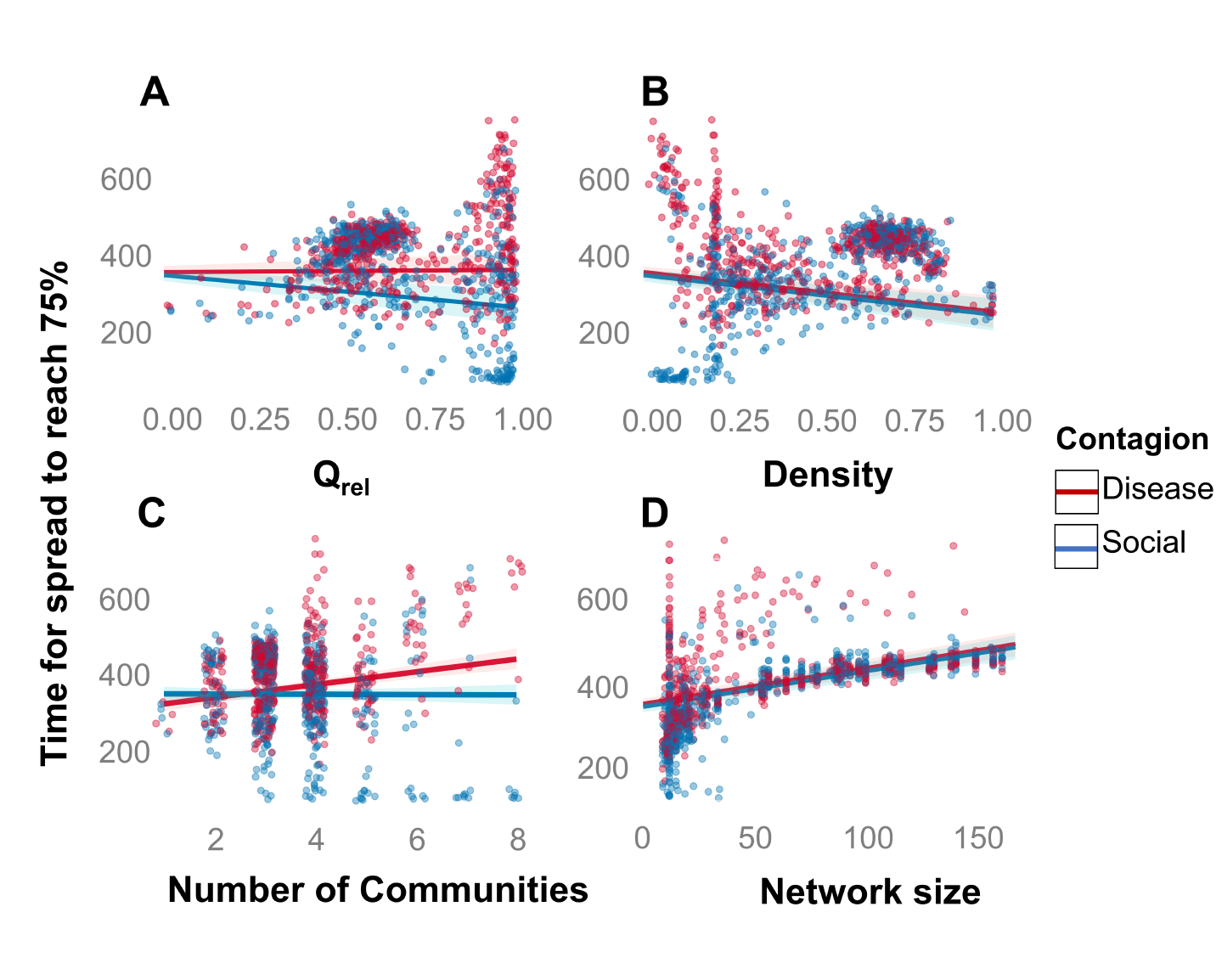
Partial effect plots illustrating how (A) modularity (Q_rel_) (B) density (C) number of communities and (D) network size influence the time for contagion spread to reach 75% of the network in disease (red) versus social (blue) contagions. Each point corresponds to the mean time for spread to reach 75% over 50 repeats in each network, and the lines represent the model’s fitted partial effects and the shaded areas depict 95% confidence intervals. Modularity, network size, and density were uncentred to their raw values for plotting. Since our regression model includes interaction effects, in (A), the number of communities is held at its mean and in (C), modularity is held at its mean.

The overall (conditional) Bayesian R^2^ of our model, which includes both fixed and random effects, was estimated at 0.323 (95% CI: 0.318, 0.328). The marginal Bayesian R^2^—reflecting only the contribution of fixed effects—was substantially lower at 0.116 (95% CI: 0.093, 0.141), suggesting that: a) in our dataset of real-world networks, network fragmentation has a relatively modest effect on contagion outcomes; and b) a substantial portion of the variance is accounted for by the random effects in the model (i.e. variation among species and networks). We calculated intra-class correlation coefficients to partition variance after accounting for fixed effects. Network-level differences explained 11.7% of the variance in spread time (95% CI: 9.9–13.7%), while species-level differences accounted for 7.8% (95% CI: 4.2–13.1%). The remaining 80.5% (95% CI: 75.5–84.1%) was residual variance, reflecting unexplained variation at the observation level beyond species, network, and fixed effects. Supplementary figure 3 illustrates how species differ in their mean contagion spread times.

## Discussion

Our results reveal that as networks become increasingly fragmented and modular, contagion spread is delayed for simple contagions (e.g., infectious disease), whereas complex contagions (e.g., information use and behaviour) can exploit dense community structure to spread to the majority of the network in less time.

Our results support and extend the theoretical findings of Evans et al. [42], which showed that highly subdivided social network architectures preferentially facilitate beneficial behavioural contagions over harmful pathogens. The effect is most distinct in networks with the highest levels of community structuring—those that are both modular and fragmented. Using networks drawn from the ASNR [45], we find that as modularity increased, the time for 75% of individuals to become infected rose for disease contagions, but only in social networks with many, small communities. In contrast, social contagions were not delayed and spread considerably faster than disease contagions when networks were fragmented and modular. However, while fragmentation and modularity in these empirical networks explains some of the differences in how these contagions play out, it accounts for only a small portion of the overall variation in outcomes, indicating that additional factors are important in shaping the dynamics of spread. This latter result demonstrates the value of using a taxonomically and behaviourally diverse sample of real-world network structures in our study. While previous research has relied on theoretical network structures [42, 41, 50], our results inherently incorporate the complexity and heterogeneity present in natural social systems to demonstrate that these theoretical patterns play a limited but potentially important role in nature.

Our data-driven modelling approach demonstrates that the slowing effect of community structure on infectious disease spread is a realistic phenomenon in natural animal societies. Our findings complement epidemiological studies showing that modular structures facilitate rapid transmission within strongly connected groups while reducing between-group spread, effectively acting as a ‘brake’ on overall transmission [38, 40, 51, 52]. By considering the interaction between network modularity and fragmentation, we extend the insights provided by [40] to show that group or sub-group size acts alongside the strength of subdivision between social units in determining epidemiological outcomes. We have focused on the overall (linear) effects of modularity and fragmentation in our analysis, but given that previous studies have provided evidence for non-linear and threshold effects for the effect of modularity on contagion outcomes [40, 53], this would be worthy of further study.

We show that the interaction between network fragmentation and contagion dynamics has important ecological and evolutionary implications, revealing how social structures can mediate trade-offs between information flow and disease spread. Although our simulations confirm that community structure and network fragmentation do interact to determine contagion outcomes, the: 1) pronounced effects being found only with high modularity and/or fragmentation; and 2) the modest effect sizes and the relatively low marginal Bayesian *R*^2^ (reflecting our fixed network structural predictors) compared to the higher overall (conditional) *R*^2^ that incorporates species and network-level variability indicate that other factors drive much of the variation in contagion spread in real-world networks. A substantial part of this unexplained variation may arise from smaller networks, where stochasticisity and sparse connections amplify the unpredictability of contagion outcomes. Additionally, individual-level heterogeneity may contribute towards the unexplained variability in our results. Future work should consider variation in node-level centrality measures to better elucidate how differences in individual network position could influence the distinct transmission dynamics of disease versus social contagions [54].

Here, we spread social information with a conformist learning rule, which has been documented across several species [30, 31, 55]. In our study, the social contagion traverses modular boundaries more efficiently, even when the same network structure impedes the spread of a simple contagion. This may be due to social reinforcement, where tight-knit subgroups with high clustering repeatedly expose individuals to the behaviour, making it easier for a complex contagion which requires multiple exposures for transmission, to spread [50, 56]. Animal social systems exhibit great diversity, with social networks varying significantly in their community structure [57, 58, 59]. Notably, our results indicate that modular networks only balance the trade-off between social and disease contagions when highly fragmented into many communities. This distinction is important because: 1) different social systems can vary considerably in the size of their groups or sub-groups (e.g., squamate reptiles can have as few as two to as many as 20,000 conspecifics in a group [60]); 2) modularity can arise at different scales—either through discrete, weakly connected groups at the population level or through hierarchical structuring within groups, as seen in multi-level societies [38, 61, 62]. In such systems, sub-group formation may help balance evolutionary pressures living in these stable social groups, and nested substructures within larger groups may influence contagion dynamics at higher social scales.

While our findings are encouraging, several modelling limitations warrant consideration. We spread social behaviour as a complex contagion, using conformist learning, yet transmission dynamics can depend on the mode of information transmission. Beck, Sheldon & Firth (2023) showed that adoption driven by the sum and strength of informed contacts favours early uptake by highly connected individuals, whereas ratio-based transmission does not [63]. Hence, alternative social-information mechanisms could yield outcomes different to our conformist model. If information spread followed simpler rules, such as proximity-triggered activity [64], network effects might resemble those of simple contagions [21]. Future work could explore social learning from high-ranking individuals or kin [65, 66] and mechanisms like payoff-based learning [67], to further unravel the differential spread of information and infection in natural systems. Moreover, we chose an intermediate transmissibility value from Evans et al. [42], where simple and complex contagions show the strongest contrast. However, high modularity mitigates contagion spread, except when contagions are transmissible enough to overcome structural barriers [40]. Accordingly, our results likely hold across parameter ranges, though structural effects may weaken under high transmissibility. Our study uses static networks, whereas many real-world social networks are dynamic; social ties frequently form and dissolve [68]. Social distancing among infected animals causes isolation [69], while individuals acquiring important information may become more central within their network [70]. Because subgroup structure and composition changes over time, static models ignore bidirectional feedback between contagion dynamics and network structure [71, 72]. Finally, our SI model provides valuable insights into the early stages of contagion uptake, but future extensions incorporating recovery (SIR) or reintroducing infected individuals into the pool (SIS) could shed light on how social structure differentially influences the long-term dynamics of simple versus complex contagions [73].

Our sample of networks from the Animal Social Network Repository (ASNR), although normalised across systems, includes diverse network types and edge definitions (e.g., spatial proximity to direct physical contact). While allowing us to generalise across animal social network types, this does introduce variability that likely added uncertainty to our conclusions [43, 74]. Namely, a limitation of our study is the inclusion of small (*<* 20 nodes) and sparse (low-density) networks, which produced greater variance in spread times and inflated Q_rel_ values due to instability in community detection [75, 76]. Although we explored this in the Supplementary Information and found that models restricted to larger or denser networks were qualitatively similar in the direction of effects, higher variance in small or sparse networks reduces precision. Because many real-world animal social networks are inherently small—often due to sampling constraints—this limitation is difficult to avoid entirely, but it should be considered when interpreting results. Another limitation of our cross-species approach is that we did not explicitly identify the species most relevant for contagion spread, given inherent differences in social learning, social tolerance, and cognitive adaptations [19, 77, 78]. More broadly, variation in sociality across species is closely linked to how groups are structured, especially in terms of group cohesion and the presence of nested or modular organisation, which likely plays a key role in shaping contagion outcomes [59]. Expanding the range of networks in the ASNR will improve the generalisability of comparative analyses and enable more networks to be classified within categories relevant for different types of contagion spread.

## Conclusions

Our study demonstrates that highly fragmented and modular networks can differentially influence contagion spread, with complex contagions propagating across subgroup boundaries more effectively than simple contagions. This differential effect has important evolutionary implications, suggesting that modular structures with many subgroups may enhance fitness of individuals that form these types of social networks by improving access to beneficial social information while mitigating the risk of acquiring infection. However, we do identify a relatively small effect size showing that these network features explain less variation in real-world systems than theoretical models, revealing the importance of data-driven modelling in translating theory practical applications in social ecology and evolution. Our findings have practical implications: for example, management interventions aimed at fragmenting social networks, such as targeted hunting, fencing, or habitat manipulation [79], may slow disease spread but will also disrupt social contagion processes, necessitating a careful assessment of unintended (behavioural) effects of these interventions. Future research should track individual-level behaviours to elucidate how individual roles vary in the spread of simple versus complex contagions, thereby clarifying the mechanisms underpinning these distinct transmission patterns and refining our understanding of the interplay between network structure, contagion dynamics, and the evolution of sociality.

## Acknowledgements

We thank the Animal Social Network Repository (ASNR) team and the many researchers who shared their data openly, making its inclusion in the repository and the present research possible.

