## Supplementary Information for "Fragmented social networks promote complex behavioural contagions over infectious disease spread"

### Modularity Calculations

Modularity (Q) was calculated following Newman's (2004) definition [1]:

$$Q = \frac{1}{2m}\sum_{i,j}\left(A_{ij} - \gamma\frac{k_i k_j}{2m}\right)\delta(c_i, c_j) \tag{1}$$

where $m$ is the number of edges, $A_{ij}$ is the weight of the $A$ adjacency matrix in row $i$ and column $j$, $k_i$ is the degree of node $i$, $k_j$ is the degree of node $j$, $c_i$ is the component of node $i$, $c_j$ is the component of node $j$, the sum goes over all pairs of nodes $i$ and $j$, and $\delta(x, y)$ equals 1 if $x = y$ and 0 otherwise.

Because the maximum modularity ($Q_{\max}$) depends on the size and number of links in the networks, direct comparisons of raw modularity values across networks can be misleading. We therefore computed relative modularity ($Q_{\text{rel}}$), by normalising observed modularity ($Q$) by its theoretical maximum ($Q_{\max}$), calculated using [2]:

$$Q_{\max} = \sum_{k=1}^{K}\left(\frac{L_k}{L}\left(1 - \frac{L_k}{L}\right)\right) \tag{2}$$

where $L_k$ is the number of edges within subgroup $k$, $L$ is the total number of edges in the network, and $K$ is the number of subgroups. Relative modularity was then defined as:

$$Q_{\text{rel}} = \frac{Q}{Q_{\max}} \tag{3}$$

### Other statistical models

To assess the robustness of our main findings, we evaluated alternative statistical models, varying assumptions about residual variance, network inclusion, and the measure of community structure. These models allow us to confirm that the observed differences between social and disease contagion dynamics are not driven by a single modelling choice.

### Heterogeneous variance model

- Heterogeneous variance model
- Standard hierarchical model
- High size–density threshold model
- Model with no *Camponotus fellah*
- Subgroup cohesion instead of $Q_{\text{rel}}$

Results from all the above models are shown in Table 1 and 2. Notably, the three-way interaction among contagion type, modularity, and number of communities remained negative and statistically clear in all five models we ran, confirming that social contagions spread more effectively in fragmented and modular networks than disease contagions do (Supplementary Figure 4).

### Standard hierachical model

We ran two main models: a heterogeneous variance model (presented in the main text) and a standard hierarchical model. The standard hierarchical model incorporated nested random effects for species within networks, but assumed a constant residual variance across all groups while the heterogenous variance model allowed residual standard deviation to vary across species and networks. Model fit was assessed using Leave-One-Out Cross-Validation (LOO). For each model, we computed the Expected Log Predictive Density (ELPD) and the LOO Information Criterion (LOOIC), where a higher ELPD and a lower LOOIC indicate better predictive performance [3]. Model comparisons were based on the difference in ELPD along with its associated standard error. The model with higher ELPD (and correspondingly lower LOOIC) was selected. Our heterogenous variance model outperformed the standard hierachical model, achieving an ELPD score of -421,829 (SE=205.5) and a LOOIC of 843,659 (SE=411.1), compared to an ELPD score of -427,135.8 (SE= 214.6) and a LOOIC of 854, 271.5 (SE= 429.2) for the standard hierachical model. Although the heterogenous variance model used more effective parameters (912.1 rather than 543.3), both models passed the Pareto k diagnostic (k < 0.7), leading us to select the heterogenous variance model. For both the heterogenous variance model ($\beta = -98.31$, 95% CI: $-105.85$, $-90.70$) and the standard hierarchical model ($\beta = -81.23$, 95% CI: $-87.65$, $-74.59$), three-way-interaction remained consistent.

### High size–density threshold model

We ran an additional model restricting analyses to networks with at least 20 nodes and density $\geq 0.15$ to assess the influence of small or sparse networks on model results. These thresholds were chosen based on the greater variance in smaller networks ($< 20$ nodes; $N = 180$, mean SE = 19.6) and sparser networks (density $< 0.15$; $N = 57$, mean SE = 40.4), compared to the more uniform variance in larger ($\geq 20$ nodes; $N = 365$, mean SE = 13.9) and denser ones (density $\geq 0.15$; $N = 488$, mean SE = 12.9) (Supplementary Information). The model specification was identical to the heterogeneous variance model presented in the main text, but applied to the reduced dataset ($n = 308$ networks). Results were qualitatively consistent with the main model in both the magnitude and direction of effects, including a negative and statistically clear three-way interaction between contagion type, modularity, and number of communities ($\beta = -38.55$, 95% CI: $[-50.72, -26.97]$). This suggests that the main conclusions are robust to the exclusion of small or sparse networks.

### Model with no *Camponotus fellah*

To test the robustness of our findings, we re-estimated the Bayesian regression model excluding ant (*Camponotus fellah*) networks, which accounted for a disproportionately large number of cases ($n = 221$) due to repeated sampling in the original study. Results were consistent with the full model (Table 2). The direction and magnitude of the three-way interaction between modularity, number of communities and contagion type ($\beta = -78.69$, 95% CI: $-86.22$, $-71.00$) supports the conclusion that social contagions consistently reach 75% of the network faster than disease contagions at high levels of both modularity and fragmentation.

### Subgroup cohesion model

We also ran a version of the model replacing $Q_{rel}$ with subgroup cohesion, defined as the proportion of contacts occurring within subgroups. Results were similar: social contagions more effectively traversed high subgroup cohesion (i.e. modularity) and a larger number of communities than disease contagions ($\beta = -69.33$, 95% CI: $-75.21$, $-63.30$).

| **Predictors** | **Heterogeneous variance** | **Standard hierarchical** | **High size-density model** |
|---|---|---|---|
| Intercept | 282.89 (261.46, 317.07) | 301.52 (276.24, 326.13) | 332.67 (294.37, 368.83) |
| Sigma Intercept | 4.80 (4.75, 4.85) | – | 4.83 (4.72, 4.93) |
| type [social contagion] | 43.38 (37.52, 49.11) | 14.07 (8.54, 19.93) | 49.08 (42.12, 56.04) |
| $Q_{rel}$ (centred) | –52.45 (–128.32, 23.19) | –31.06 (–95.01, 36.49) | –15.29 (–147.99, 116.73) |
| Number of communities | 16.73 (10.31, 23.47) | 14.40 (8.77, 20.11) | 9.25 (2.56, 15.71) |
| Density (centred) | –102.69 (–141.82, –64.76) | –87.67 (–122.67, –52.89) | –131.14 (–183.03, –80.05) |
| Network size (centred) | 0.87 (0.71, 1.03) | 0.99 (0.85, 1.15) | 0.80 (0.66, 0.94) |
| type × $Q_{rel}$ | 12.44 (–49.37, 24.29) | 13.60 (–39.30, 15.87) | –10.60 (–59.50, 39.21) |
| type × # communities | –17.06 (–18.80, –15.33) | –12.91 (–14.58, –11.30) | –17.56 (–19.58, –15.61) |
| $Q_{rel}$ × # communities | 19.44 (2.28, 40.98) | 12.95 (–6.32, 31.38) | –2.36 (–32.52, 29.23) |
| type × $Q_{rel}$ × # communities | –98.31 (–105.85, –90.70) | –81.23 (–87.65, –74.59) | –38.55 (–50.72, –26.97) |
| *Bayesian $R^2$ (marg.)* | 0.323 (0.318, 0.328) | 0.372 (0.369, 0.376) | 0.200 (0.191, 0.210) |
| *Residual variance (prop.)* | 0.805 (0.755, 0.841) | • | 0.738 (0.628, 0.825) |
| *Proportion variance (random effects)* | | | |
| Species | 0.078 (0.042, 0.131) | • | 0.212 (0.126, 0.330) |
| Network | 0.117 (0.099, 0.137) | • | 0.050 (0.035, 0.068) |

Supplementary Table 1. Posterior estimates (95% credible intervals) for the heterogeneous variance model, the standard hierarchical model, and the model restricted to large, dense networks. "•" indicates not applicable.

| **Predictors** | **No-ants model** | **Subgroup cohesion** |
|---|---|---|
| Intercept | 348.96 (300.67, 398.14) | 280.11 (254.22, 304.87) |
| Sigma Intercept | 4.85 (4.81, 4.89) | 4.78 (4.73, 4.83) |
| type [social contagion] | –57.72 (–65.67, –49.81) | 66.93 (62.13, 72.01) |
| $Q_{rel}$ (centred) | 22.74 (–82.83, 126.70) | – |
| Number of communities | 10.30 (0.21, 20.89) | 20.40 (15.19, 25.69) |
| Density (centred) | –116.27 (–171.83, –65.37) | –90.33 (–122.93, –55.78) |
| Network size (centred) | 1.22 (0.76, 1.67) | 0.91 (0.75, 1.08) |
| type × $Q_{rel}$ | 16.50 (–40.38, 25.43) | – |
| type × # communities | –2.37 (–4.49, –0.29) | –27.38 (–28.91, –25.97) |
| $Q_{rel}$ × # communities | –9.00 (–40.15, 20.98) | – |
| type × $Q_{rel}$ × # communities | –78.69 (–86.22, –71.00) | – |
| *Bayesian $R^2$ (marg.)* | 0.260 (0.254, 0.267) | 0.334 (0.330, 0.339) |
| *Residual variance (prop.)* | 0.759 (0.714, 0.797) | 0.800 (0.751, 0.839) |
| *Proportion variance (random effects)* | | |
| Species | 0.048 (0.018, 0.093) | 0.082 (0.044, 0.134) |
| Network | 0.193 (0.163, 0.226) | 0.118 (0.100, 0.138) |

Supplementary Table 2. Posterior estimates (95% credible intervals) for the model excluding ant networks and the subgroup cohesion model.

### Collinearity diagnosis

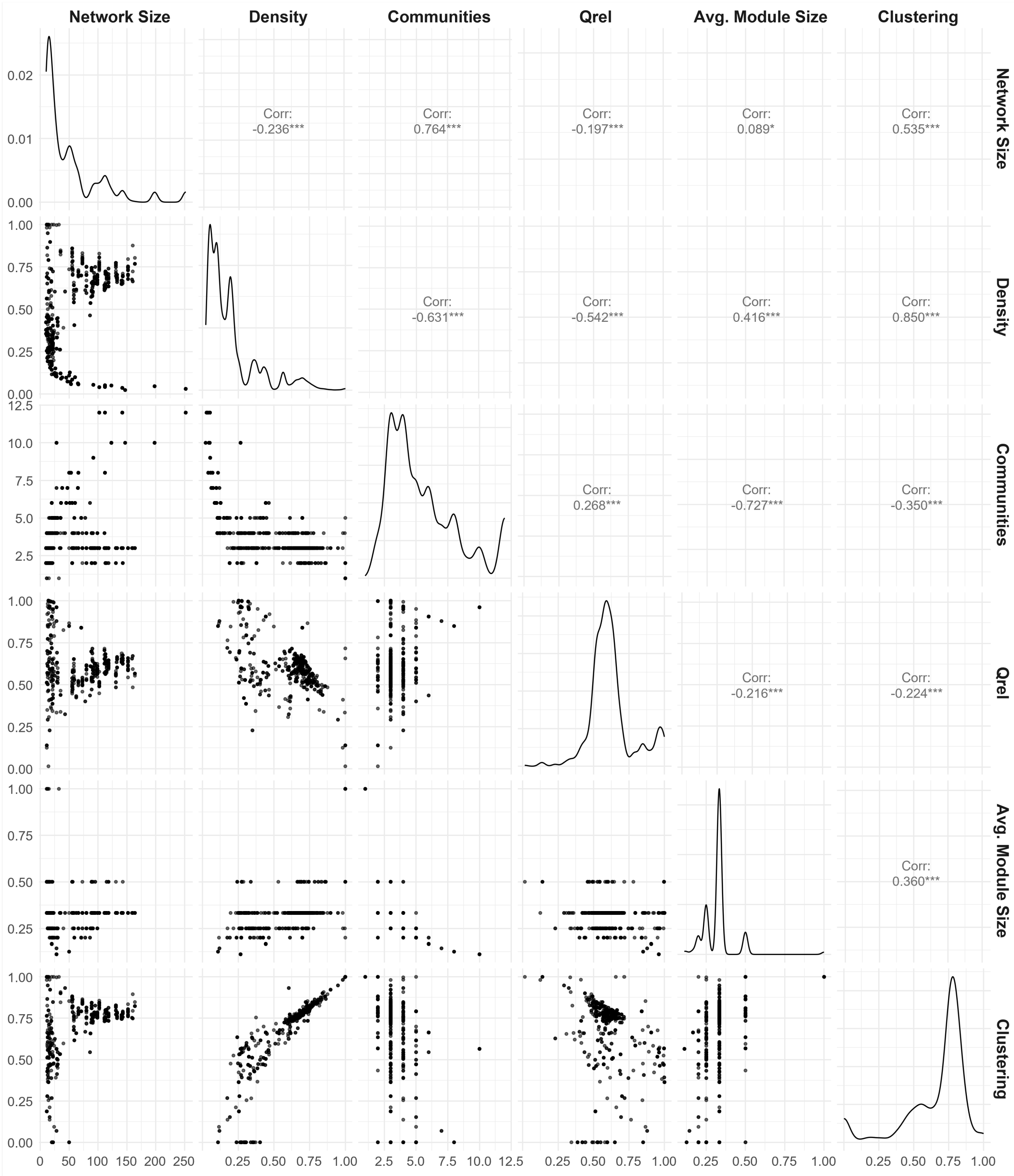


Supplementary Figure 1. Pairwise relationships among network metrics. Pearson correlation coefficients are shown in the upper panels, scatterplots in the lower panels, and metric distributions on the diagonal. Variables with pairwise correlations above 0.7—clustering coefficient and average module size—were excluded from downstream analyses due to collinearity.

### Species-level random effects

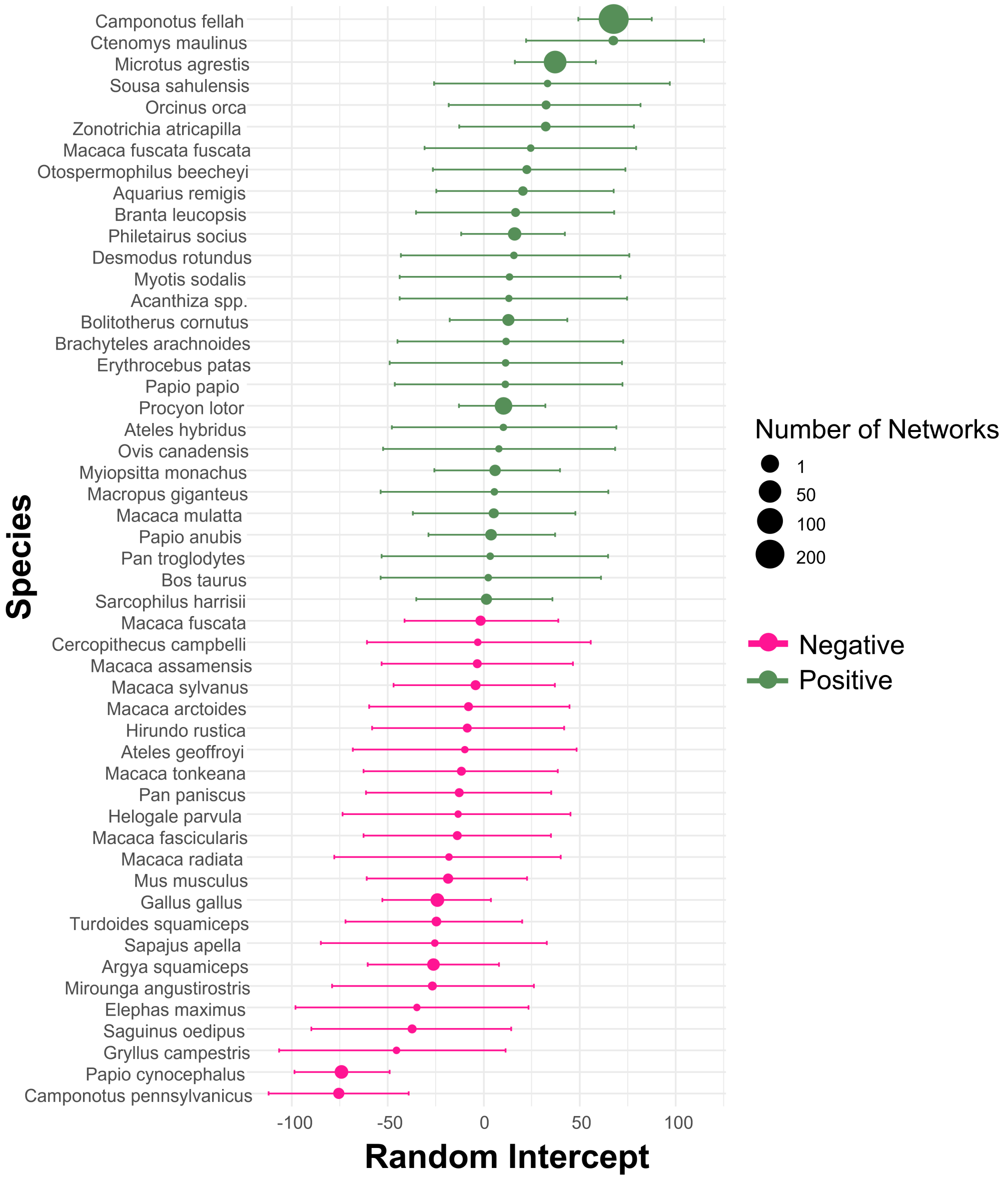


Supplementary Figure 2. Species-level random intercepts from the Bayesian model, illustrating variation among species in the baseline time for contagion spread. Circles show each species' posterior mean intercept, and horizontal lines represent 95% credible intervals. Circle size is proportional to the number of networks contributed by that species, and circle colour indicates whether the random intercept is positive (green) or negative (pink) relative to zero.

### Posterior predictive model check

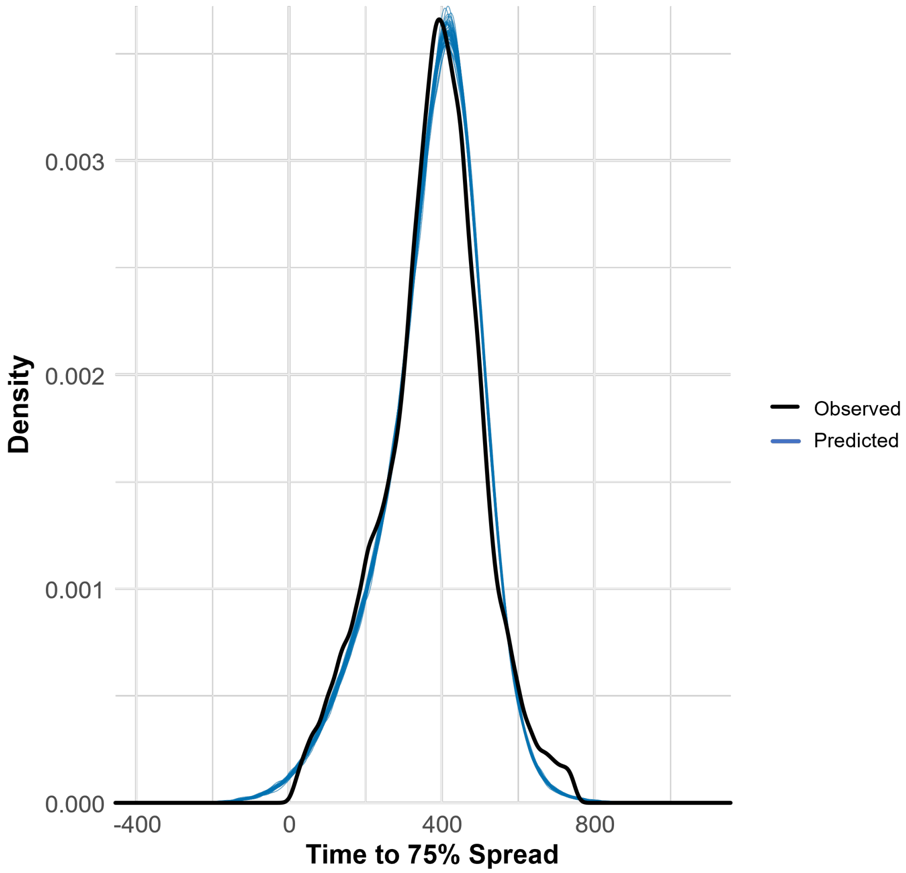


Supplementary Figure 3. Posterior predictive check using a density overlay: the black curve shows the observed distribution of time to 75% spread, and the blue curves represent 30 posterior predictive draws from the Bayesian model, demonstrating model fit.

### Three-way interaction plots of all supplementary models

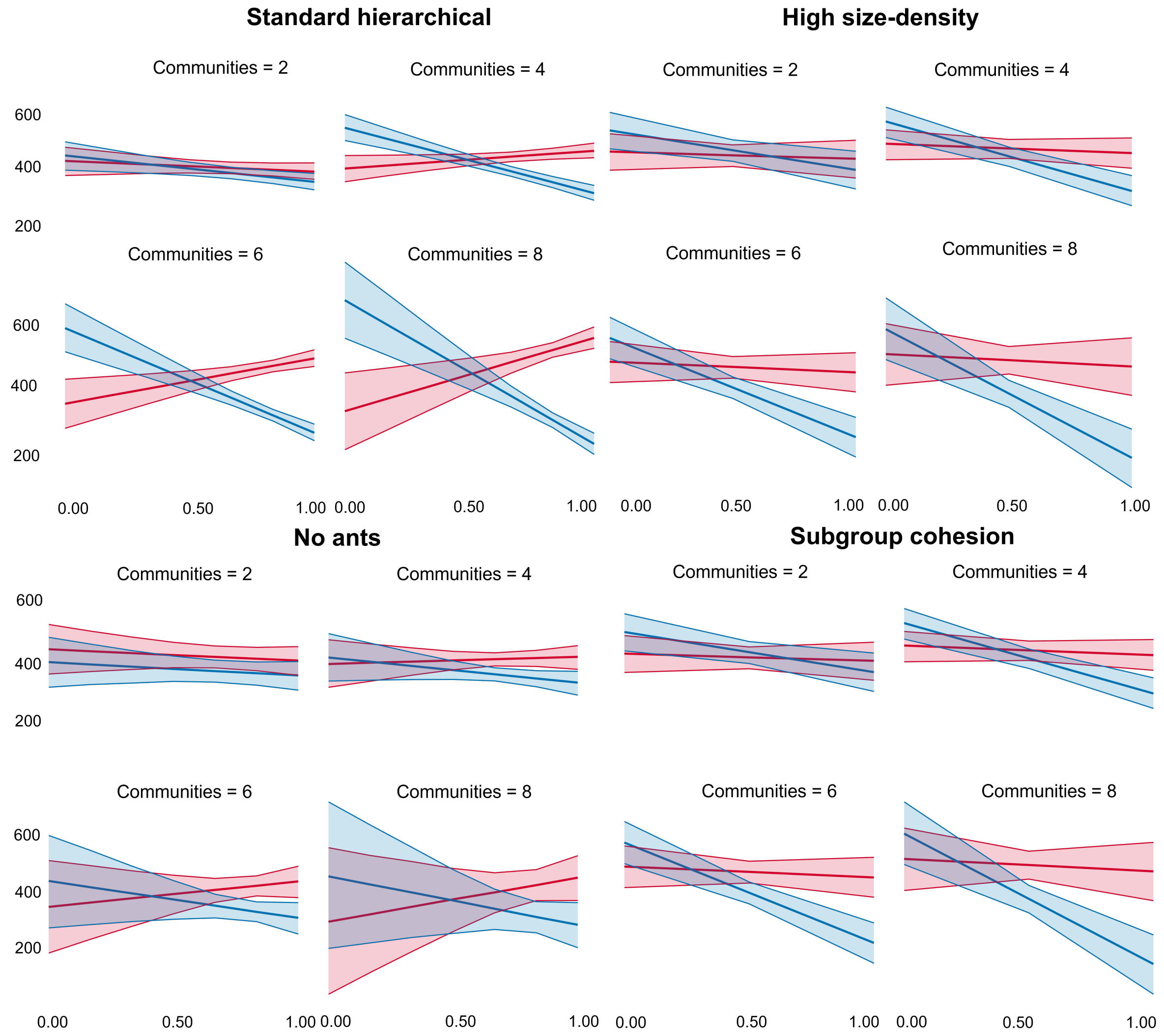


Supplementary Figure 4. Three-way interaction plots from all supplementary models described in the text. Each panel shows the interaction between modularity ($Q_{rel}$) and number of communities on the time to 75% contagion spread. Results are shown for the standard hierarchical model, high size–density model, model excluding *Camponotus fellah* ("No ants"), and the model using subgroup cohesion in place of relative modularity. Red lines indicate simple contagions (infection), while blue lines indicate complex contagions (information). Shaded areas represent 95% credible intervals.
